# SPECT/CT imaging, biodistribution and radiation dosimetry of a ^177^Lu-DOTA-integrin αvβ6 cystine knot peptide in a pancreatic cancer xenograft model

**DOI:** 10.1101/2021.03.23.436654

**Authors:** Sachindra Sachindra, Teresa Hellberg, Samantha Exner, Sonal Prasad, Nicola Beindorff, Stephan Rogalla, Richard Kimura, Sanjiv Sam Gambhir, Bertram Wiedenmann, Carsten Grötzinger

## Abstract

**Introduction:** Pancreatic ductal adenocarcinoma (PDAC) is one of the most aggressive malignant neoplasms, as many cases go undetected until they reach an advanced stage. Integrin αvβ6 is a cell surface receptor overexpressed in PDAC. Consequently, it may serve as a target for the development of probes for imaging diagnosis and radioligand therapy. Engineered cystine knottin peptides specific for integrin αvβ6 have recently been developed showing high affinity and stability. This study aimed to evaluate an integrin αvβ6-specific knottin molecular probe containing the therapeutic radionuclide ^177^Lu for targeting of PDAC.

**Methods:** The expression of integrin αvβ6 in PDAC cell lines BxPC3 and Capan2 was analyzed using RT-qPCR and immunofluorescence. In vitro competition and saturation radioligand binding assays were performed to calculate the binding affinity of the DOTA-coupled tracer loaded with and without lutetium to BxPC3 and Capan2 cell lines. To evaluate tracer accumulation in the tumor and organs, SPECT/CT, biodistribution and dosimetry projections were carried out using a Capan2 xenograft tumor mouse model.

**Results:** RT-qPCR and immunofluorescence results showed high expression of integrin αvβ6 in BxPC3 and Capan2 cells. A competition binding assay revealed high affinity of the tracer with IC_50_ values of 1.69 nM and 9.46 nM for BxPC3 and Capan2, respectively. SPECT/CT and biodistribution analysis of the conjugate ^177^Lu-DOTA-integrin αvβ6 knottin demonstrated accumulation in Capan2 xenograft tumors (3.13 ± 0.63 %IA/g at day 1 post injection) with kidney uptake at 19.2 ± 2.5 %IA/g, declining much more rapidly than in tumors.

**Conclusion:** ^177^Lu-DOTA-integrin αvβ6 knottin was found to be a high-affinity tracer for PDAC tumors with considerable tumor accumulation and moderate, rapidly declining kidney uptake. These promising results warrant a preclinical treatment study to establish therapeutic efficacy.

## Introduction

Pancreatic ductal adenocarcinoma (PDAC) is one of the most aggressive malignant neoplasms and accounts for 80-90% of all pancreatic cancer cases. Although the incidence of PDAC is low, in cancer-related deaths, it ranks seventh globally (*1*). Owing to its poor prognosis, the 5-year survival rate is 9% with merely 24% of patients surviving for a year (*2*). This is mainly because PDAC patients rarely exhibit symptoms before an advanced stage of the disease has been reached, and due to the lack of appropriate diagnostic and therapeutic options. Surgical resection of the tumor, part of the pancreas, and other nearby digestive tract organs remains to be the only curative treatment for early-stage PDAC patients. Gemcitabine has been used for several years as a baseline chemotherapeutic treatment. Lately, combination therapy of gemcitabine with folfirinox and nab-paclitaxel demonstrated improved results in comparison to the use of gemcitabine alone (*3,4*). However, high resistance of PDAC to chemotherapy dilutes its efficacy (*5,6*).

Integrins are heterodimeric transmembrane cell surface proteins that mediate cell-to-cell and cell-to-extracellular matrix (ECM) adhesion (*7,8*). Many integrins, including integrin αvβ6, were reported to be upregulated in various cancers such as breast cancer, gastric cancer, colorectal cancer, lung cancer, ovarian cancer and PDAC (*9*–*14*). In well-differentiated PDACs, integrin αvβ6 overexpression was identified in 100% of the samples (*13,14*). Moreover, integrin αvβ6 overexpression has been recognized as a prognostic marker for reduced survival in non-small cell lung cancer (*15*), gastric carcinoma (*16*), colorectal cancer (*17*) cervical squamous cell carcinoma (*18*) and PDAC (*19*). Remarkably, integrin αvβ6 expression was found to be higher in PDAC than in chronic pancreatitis (*20*). These findings support the utilization of integrin αvβ6 as a target for the development of new diagnostic and therapeutic tools.

Cystine knot peptides (knottins) represent small peptides of approximately 4 kDa with three threaded disulfide bonds that form a topological knot. Such a structural motif is known as a cystine knot (*21,22*). One of the advantages of knottins is the high variability of backbone residues that may be used to modulate tumor and kidney uptake (*23*). Previously, we have developed the optimized knottin R_0_1-MG that shows low single-digit nanomolar binding affinity for integrin αvβ6 (*23,24*). Recently, the first clinical study with R_0_1-MG demonstrated clinical potential for targeting PDAC in humans (*24*).

In the current imaging, biodistribution and dosimetry study, a lutetium-177 DOTA conjugate of this knottin was evaluated as a candidate for therapeutic purposes in a PDAC xenograft model. Binding affinity of the ^177^Lu tracer was found to be in the low single-digit nanomolar range. SPECT/CT imaging and biodistribution of the knottin tracer ^177^Lu-DOTA-integrin αvβ6 revealed substantial accumulation of the tracer in the tumor as well as faster renal clearance. Our studies demonstrate the potential of the ^177^Lu integrin αvβ6 knottin as a therapeutics candidate in PDAC.

## Material and Methods

### Cell culture

The human pancreatic adenocarcinoma cell lines BxPC3 and Capan2 were obtained from ATCC/LGC Standards (Wesel, Germany). They were cultured in RPMI 1640 and McCoy’s 5A (Modified) Medium (both Biochrom, Berlin, Germany), respectively, and supplemented with 10% fetal calf serum (Biochrom, Berlin, Germany) for cultivation in a humidified atmosphere at 37 °C with 5% CO_2_.

### Reverse transcription quantitative real-time PCR (RT-qPCR)

Total RNA was isolated from cell lines or kidney and liver of mouse using the RNeasy Mini Kit (Qiagen, Hilden, Germany) according to the manufacturer’s protocol. RNA was treated with DNase I (Sigma Aldrich, Munich, Germany) before reverse transcription with the High Capacity cDNA Reverse Transcription Kit (Applied Biosystems, Waltham, MA, USA). Quantitative real-time PCR was performed with Blue S’Green qPCR 2X mix (Biozym Scientific GmbH, Oldendorf, Germany), 0.5 µM primer and 20 ng cDNA in a total reaction volume of 10 µL on a Bio-Rad CFX96 Real-Time-System. PCR settings were 95 °C for 30 sec, followed by 45 cycles of 95 °C for 5 sec and 61 °C for 30 sec. All primers were designed using NCBI Primer-BLAST software. They were manufactured by Tib-MolBiol (Berlin, Germany) and their sequences are indicated below. Plotted values were normalized on the geometric mean of UBC and GAPDH (for cells and xenografts) or GAPDH only (for mouse kidney and liver) using the ^ΔΔ^C_t_ method.

hITGB6-Fwd: 5’-ACTGCCTGCTTATTGGACCTC-3’

hITGB6-Rev: 5’-ATCACACCTTTCGCCAACTC-3’

hUBC-Fwd: 5’-ATTTGGGTCGCAGTTCTTG-3’

hUBC-Rev: 5’-TGCCTTGACATTCTCGATGGT-3’

hGAPDH-Fwd: 5’-TGCACCACCAACTGCTTAGC-3’

hGAPDH-Rev: 5’-GGCATGGACTGTGGTCATGAG-3’

mITGB6-Fwd: 5’-CTCACgggTACAgTAACgCAT-3’

mITGB6-Rev: 5’-AATgAgCTCTCAggCAggCT-3’

### Immunofluorescence

For immunofluorescent staining of tumor tissue, cells grown on glass coverslips were fixed with 1:1 methanol/acetone for two minutes and air-dried. After washing with PBS (Biochrom, Berlin, Germany) and blocking with 5% goat serum in PBS for 30 minutes, coverslips were incubated with a rabbit IgG against human integrin β6 (#HPA023626, Atlas Antibodies, Bromma, Sweden) diluted in 0.1% BSA in PBS in a wet chamber for one hour at room temperature. After washing, coverslips were incubated with the secondary antibody goat-anti-rabbit-Cy3 (Jackson ImmunoResearch, West Grove, USA; 2.5 µg/mL diluted in 0.1% BSA in PBS) for 30 minutes. After washing with PBS, nuclei were stained with 1 µg/mL TOTO-3 (Invitrogen) in PBS for 5 minutes. Finally, the cells were fixed with 96% ethanol for two minutes, embedded with Immu-Mount (Thermo Fisher Scientific, Waltham, USA) and analyzed using a confocal laser-scanning microscope (LSM510, Carl Zeiss, Jena, Germany).

### Integrin-αvβ6 peptide radiolabelling (iodination)

The previously described cystine knot peptide R_0_1-MG (GCILNMRTDLGTLLFR-CRRDSDCPGACICRGNGYCG-DOTA) specific for integrin αvβ6 was utilized in this study. The DOTA-knottin was labelled with ^125^I using the chloramine-T method. Chloramine-T and sodium metabisulfite solutions were freshly prepared in water. For labelling, 10 µl of DOTA-knottin (stock 1 mM) with 15 µl of sodium phosphate buffer (0.5 M, pH 7.6) was mixed with 37 MBq carrier-free Na^125^I (NEZ033L010MC, Perkin Elmer, Waltham, US). 4 μl chloramine T (1 mg/ml) were added to start the reaction and after 30 seconds, 4 μl sodium metabisulfite (2 mg/ml) were added to stop the iodination. Labelled radioactive peptide was separated from unlabeled peptide by HPLC purification (Agilent ZORBAX 300 Extend-C18 column) using a gradient from 20-50% acetonitrile (+0.1 % TFA) against water (+0.1 % TFA) for 20 min. To determine the retention time of the radioactive peptide, 1-2 µl of the reaction mixture were analyzed before the purification run. The fraction containing the radiolabeled peptide peak was then collected, diluted with binding buffer (50 mM HEPES pH 7.4, 5 mM MgCl_2_, 1 mM CaCl_2_, 0.5 % BSA, cOmplete protease inhibitors) to prevent radioautolysis, aliquoted and stored at −80 °C.

### Non-radioactive labelling with ^nat^Lu

Non-radioactive complexes of DOTA-knottin with native lutetium was generated by incubation of 30 µM peptide conjugate dissolved in 5 µl buffer (sodium acetate/acetic acid buffer, 0.5 M, pH 5.4) using a 30-fold molar excess of the Lu^3+^ ion (^nat^Lu-DOTA-knottin) or no metal ion (control DOTA-knottin). The reaction volume was made up to 30 µl with water. The reaction was carried out for 10 minutes at 80°C.

### Saturation binding assay

To determine the dissociation constant (K_d_) and the maximum number of binding sites (B_max_) of the radioligand saturation binding assay was performed. For this, approximately 50,000 cell per well were seeded in a 96 well plate and incubated overnight at 37 °C. Cells were then incubated with binding buffer and varying concentration (0-20 nM) of labeled peptide either with (non-specific binding) or without 1 µM of unlabeled peptide (total binding). Non-specific binding was subtracted from total binding to obtain specific binding. All three datasets were plotted with GraphPad Prism and fitted using nonlinear regression (one site - total and non-specific, one site - specific binding). The software provides Bmax in the same value as the respective y-axis, in this case cpm. The following calculations were performed to obtain B_max_ in receptor sites per cell. In the first step, the specific activity of ^125^I (2175 Ci/mmol) was transformed into dpm/mmol by multiplication with 2.22*10^12^. This was multiplied with the counter efficiency of 50% to get cpm/mmol and subsequently converted to cpm/fmol:

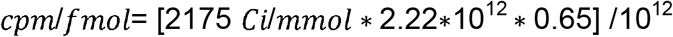

In a second step, the B_max_ value, calculated by the software in cpm, was divided by cpm/fmol to derive the amount of substance in fmol. This was multiplied with the Avogadro constant (6.02*10^8^/fmol) to obtain the number of molecules before division by seeded cells (50,000 in this case):

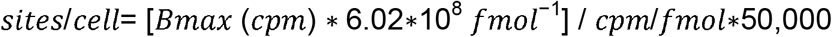

### Competition binding assay

For competitive radioligand binding assay, approximately 50,000 cells per well were seeded in a 96 well plate incubated overnight at 37°C. Next day, cells were incubated in binding buffer containing 100,000 cpm (counts per minute) ^125^I DOTA-knottin and increasing concentrations of unlabeled peptide. After 2 hours of incubation, cells were washed for 2-3 times with ice-cold washing buffer (50 mM Tris-HCl, pH 7.4, 125 mM NaCl, 0.05% BSA) and lysed with 1 N NaOH (80 µl/well). The lysed cells were transferred to vials and measured in a gamma counter (Wallac 1470 Wizard, Perkin Elmer, Waltham, MA, USA). Obtained cpm values were plotted with GraphPad Prism and the data fitted using nonlinear regression (one site - fit Ki, one site - fit logIC_50_).

### Cell irradiation

Cells were irradiated using a GSR D1 gamma irradiator (Gamma-Service Medical GmbH, Leipzig, Germany) with a ^137^cesium source. The cells with the respective culture medium were placed inside the radiation chamber and required dose was achieved by adjusting the tray level and duration of the exposure. After irradiation, the cell lines were incubated at 37 °C without medium change for the appropriate duration.

### Cell count (DAPI staining)

Cells were seeded at a density of 5,000 cells per well in a 96 well plate and incubated overnight at 37°C. After irradiating with 0, 0.0, 0.2, 2, 4, 6, 8, 10, 15 or 30 Gy, cells were incubated at 37 °C for 96 hours. Cells were then washed with PBS and fixed with 4% PFA for 10 min at room temperature. Thereafter, cells were stained with DAPI (1:5,000 in PBS/0.1% Tx-100) and incubate at room temperature for 10 min. Before image acquisition, wells were filled with 80 µL of PBS and images were acquired using an automated microscope (IN Cell Analyzer 1000; GE, Reading, UK) with a 4x objective. Image stacks were analyzed and nuclei were counted by the IN Cell software.

### Clonogenic assay

Cells were seeded at a density of 5,000 cells per well in a 12 well plate and incubated overnight at 37 °C. After irradiation with 0, 2, 4, 6 or 8 Gy, cells were incubated at 37 °C without medium change for 7 days. Cells were then washed with PBS and fixed with 70% ethanol for 10 min, stained with 0.2% crystal violet solution for another 10 min and carefully rinsed with tap water. Plates were dried overnight and digitized using an Odyssey infrared scanner (700 nm channel, intensity 3, 84 μm resolution and medium quality). For quantification, images were analyzed using the Colony Area plugin for ImageJ.

### Xenografts

For *in vivo* experiments, at least 8-week-old female athymic NMRI*-*Foxn1^nu^ /Foxn1^nu^ mice (Janvier Labs, Saint-Berthevin, France) were used. Animal care followed institutional guidelines and all experiments were approved by local animal research authorities. For the generation of tumor xenografts, 5 × 10^6^ cells of Capan2 cells were inoculated subcutaneously into the left and right shoulder (1:1 phosphate-buffered saline [PBS]/Matrigel Basement Membrane Matrix High Concentration, Corning, Corning, USA). Tumors were allowed to grow for two to four weeks (tumor volume > 100 mm^3^) after cell inoculation.

### Radiochemical labeling with lutetium-177 (^177^Lu)

Radiolabeling of the DOTA-knottin was carried out manually using ^177^Lu-LuCl_3_ and a reagent kit from ITG Isotope Technologies Garching GmbH (Garching, Germany). A total of 40 µL (stock solution of peptide at 1 µg/µL in 10% DMSO in water) was added to 500 µL of ascorbate buffer solution prepared from the kit and the resulting mixture was then added to 35 µL of ^177^Lu-LuCl_3_ (2 GBq in aqueous 0.04 M HCl) and the reaction mixture was heated at 80 °C for 35 minutes followed by cooling for 10 minutes at room temperature. The product was then diluted with 0.5 mL of saline and the pH was adjusted to 7 using NaHCO_3_ prior to injection in mice. Radiochemical purity determined by Radio-HPLC was found to be higher than 98% in all cases and the product had a molar activity of approximately 44 GBq/µmol.

### SPECT/CT imaging

SPECT and CT imaging were performed using the nanoSPECT/CTplus scanner (Mediso, Hungary /Bioscan, France). Mice were anesthetized using 1-2% isoflurane with oxygen at a flow rate of approximately 0.5 l/min. After a low-dose CT scan for positioning the tumors and kidneys in the scan range the SPECT acquisition was started directly before intravenous injection of approximately 50 MBq of ^177^Lu-integrin αvβ6-DOTA (0.1-0.15 ml).

Nine consecutive multi-pinhole SPECT images of 10 min duration each (5 angular steps a 60 sec, 2 bed positions) were acquired. Additional individual scans of 35-60 min duration (5 angular steps á 60-180 sec, 2 bed positions) were performed up to 8 days to assess biodistribution kinetics.

### Biodistribution studies

Tumor-bearing mice were injected with approximately 50 MBq of ^68^Ga-labeled DOTA-peptide or 1 MBq of ^177^Lu-labeled DOTA-peptide to the tail vein via a catheter. Mice were sacrificed and dissected one, two, three and eight days post injection. Tumor, blood, stomach, pancreas, small intestine, colon, liver, spleen, kidney, heart, lung, muscle and femur samples were weighed and uptake of radioactivity was measured by a gamma counter. To determine the effect of unlabeled ligand on tumor uptake, 200 nmol CG34 peptide (100-fold excess) was co-injected.

### Dosimetry of tissue and tumor

Assuming that the organ-to-whole-body activity concentration ratio in mice would equal that in humans, the injected activity concentration (%IA/g) acquired from the mouse biodistribution study was transformed to human whole-organ percentage of injected activity per gram of tissue (%IA/g) _human._ The mouse uptake data were extrapolated to humans by relative scaling of mass and time using the following two equations,

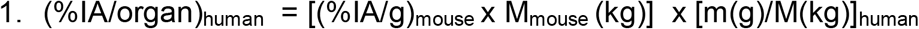

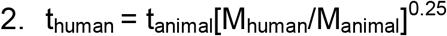

where %IA/g is percentage of injected activity per gram of tissue, %IA/organ is percentage of injected activity per organ; m is organ mass of mouse or human, M is body mass of mouse or human; t is time. The absorbed doses per activity were calculated by using the extrapolated human source organ residence times as input in the OLINDA/EXM 1.1 software (Organ Level Internal Dose Assessment Code, Vanderbilt University, Nashville, USA), with the reference adult male (*25*).

### Statistical analysis and data availability

All statistical analyses were performed using GraphPad Prism 5.04. EC_50_/IC_50_ values were determined by nonlinear sigmoidal curve fitting with variable slope setting. All presented data are based on independent experiments. Numerical data for this study have been deposited in an open data repository for public access: http://doi.org/10.5281/zenodo.4362503

## Results

### Target expression and tracer affinity

As integrin β6 forms heterodimers only with integrin αv, detection of the β6 subunit mRNA or protein will be informative about the dimer, too. To identify the optimal animal model for imaging, integrin β6 mRNA was measured by RT-qPCR in human PDAC cell lines, corresponding xenografts, A549 lung adenocarcinoma and HT29 colorectal adenocarcinoma cells as well as mouse kidney and liver. Capan2 and BxPC3 cells and xenografts showed the highest abundance of integrin β6 mRNA (Figure 1A). Mouse kidney and liver showed around 1,000-fold less integrin β6 mRNA, indicating a significantly lower expression of the corresponding mouse receptor in these organs. Immunofluorescence staining detected integrin β6 on the plasma membrane of Capan2 cells (Figure 1B).

**Figure 1:**
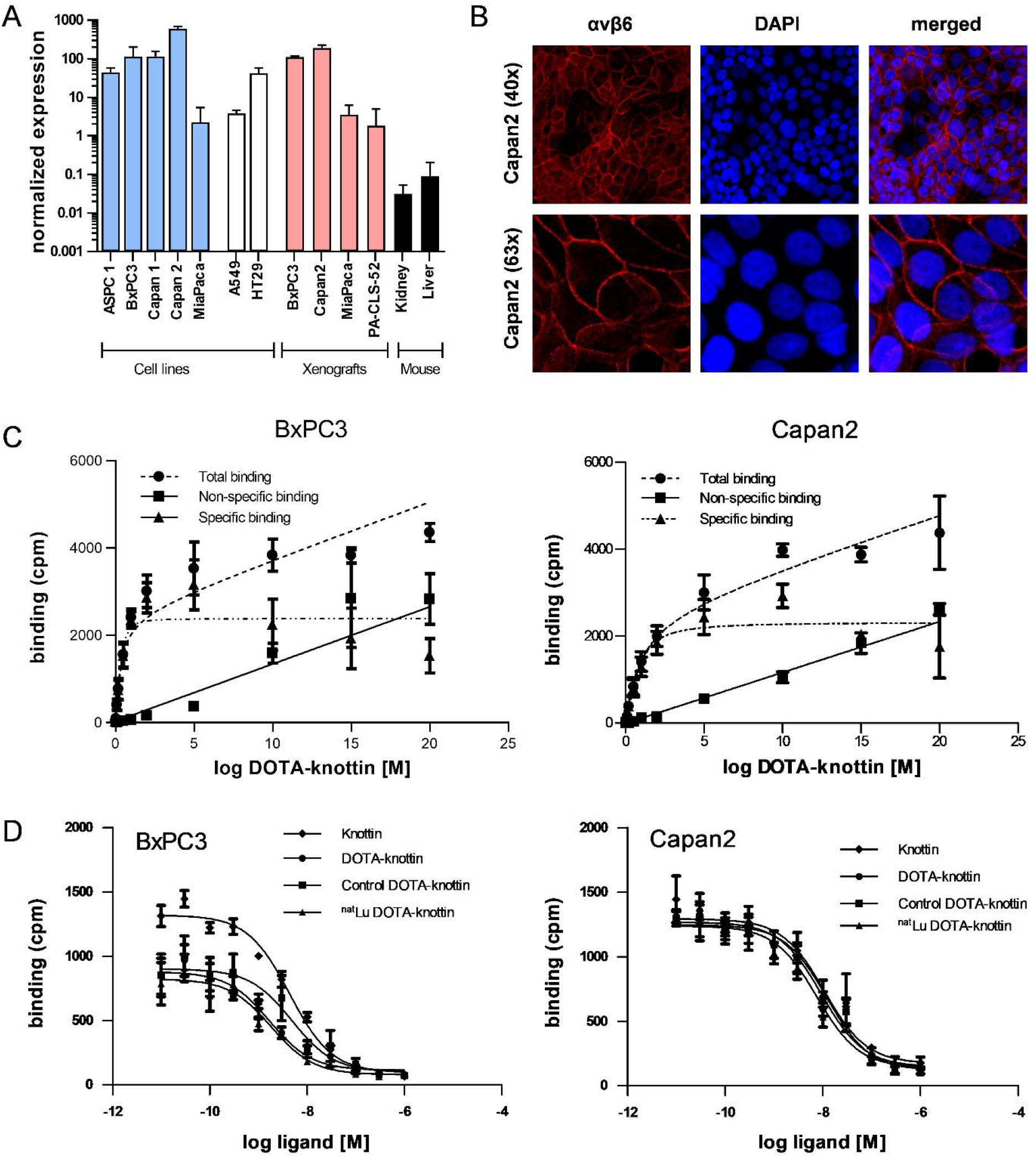
Integrin αvβ6 mRNA, protein localization, binding sites and tracer affinity. (A) Integrin αvβ6 mRNA was measured by RT-qPCR in human cell lines of different origin, xenografts and mouse kidney and liver. (B) Fluorescence images show staining of integrin αvβ6 (antibody, red) and nucleus (TOTO-3, blue) in Capan2 cells. (C) Dissociation constant (K_d_) and number of binding sites per cell were determined in BxPC3 and Capan2 cells using a saturation binding assay. (D) A competition binding assay was performed to derive the inhibitory constant (K_i_) of DOTA-knottin in both cell lines. The binding curves show tracer displacement by increasing concentrations of unlabeled knottin, DOTA-knottin, control-DOTA-knottin and ^nat^Lu-DOTA-knottin. For the latter two, DOTA-knottin was either complexed with native lutetium (^nat^Lu-DOTA-knottin) or treated the same but without the metal ion (control-DOTA-knottin). Data represent mean ± S.E.M (3 ≤ n ≤ 4).

To determine the binding affinity of the integrin αvβ6-specific DOTA-knottin to its target, saturation and competition binding assays on BxPC3 and Capan2 cells were performed (Figure 1C, D; Table 1). Saturation binding assays were performed to determine dissociation constant (K_d_) and maximum number of binding sites per cell (B_max_). K_d_ values for BxPC3 and Capan2 cells were found to be 0.30 nM and 0.75 nM, respectively. Correspondingly, the B_max_ values for BxPC3 and Capan2 were shown to be approximately 11,800 and 11,500 binding sites/cell. ^125^I-labeled DOTA-knottin showed binding to both cell lines, BxPC3 and Capan2, displaced by the unlabeled peptide in a concentration-dependent manner. The inhibitory constant (K_i_) values of DOTA-knottin for BxPC3 and Capan2 were calculated to be 1.69 nM and 9.46 nM respectively. To determine the effect of lutetium chelation on the probe’s affinity for integrin αvβ6, binding of ^nat^Lu-DOTA-knottin and control-DOTA-knottin (without ^nat^Lu) on BxPC3 and Capan2 was examined. The complexation of the ^nat^Lu ion did not compromise the high affinity of the tracer (Table 1).

**Table 1:** Saturation and competition binding assay results for BxPC3 and Capan2 cells. Data are presented as mean ± S.E.M. (3 ≤ n ≤ 4)

|  | Kd (nM) | Bmax<br>(Sites/cell) | Ki (nM) |  |  |  |
| --- | --- | --- | --- | --- | --- | --- |
|  |  |  | knottin | DOTA<br>knottin | control DOTA<br>knottin | <sup>natLu</sup> DOTA<br>knottin |
| **BxPC3** | 0.30 ± 0.08 | 11,874 ± 897 | 4.36 ± 0.16 | 1.69 ± 1.3 | 4.85 ± 1.94 | 1.82 ± 1.39 |
| **Capan2** | 0.75 ± 0.19 | 11,545 ± 887 | 10.79 ± 4.03 | 9.46 ± 5.21 | 12.06 ± 2.87 | 7.54 ± 1.27 |

### Radiosensitivity of BxPC3 and Capan2 cells

To evaluate the suitability of BxPC3 and Capan2 cells as model cell lines in terms of their radiosensitivity, two approaches were taken. Both cell lines were irradiated with different doses (0, 0.05, 0.2, 2, 4, 6, 8, 10, 15 and 30 Gy) of gamma radiation from a ^137^Cs source. After 96 hours of incubation, nuclei were stained and counted. The IC_50_ values for the resulting growth inhibition/cell death in BxPC3 and Capan2 cells were found to be 4.3 Gy and 5.5 Gy, respectively (Figure 2A). To evaluate radiation effects on colony formation, both cell lines were treated with different doses (0, 2, 4, 6 and 8 Gy) of radiation. After 7-8 days of incubation, colonies were fixed, stained and counted (Figure 2B, C). BxPC3 cells (IC_50_ 1.3 Gy) appeared to be slightly more radiation-sensitive compared to Capan2 (IC_50_ 2.2 Gy) in this assay.

**Figure 2:**
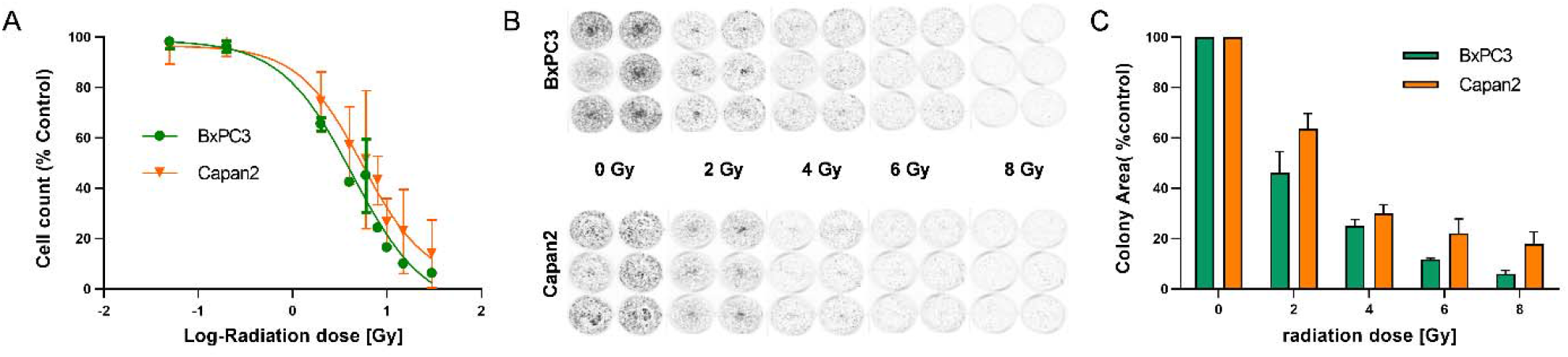
Radiosensitivity of BxPC3 and Capan2 cells. (A) BxPC3 and Capan2 cells were exposed to different doses of radiation (0, 0.05, 0.2, 2, 4, 6, 8, 10, 15 and 30 Gy). 96 hours later, nuclei were stained and counted. (B, C) BxPC3 and Capan2 cells were irradiated with 0, 2, 4, 6 or 8 Gy. Cells were incubated for 1 week. Colonies were stained and colony area was measured. Data represent mean ± S.E.M. (n=5).

### SPECT imaging in a Capan2 xenograft mouse model of pancreatic cancer

While the integrin αvβ6-specific knottin and its DOTA conjugate previously had been used for tumor imaging studies employing ^18^F, ^64^Cu or ^99m^Tc, no data regarding the biodistribution of the ^177^Lu-labeled DOTA-knottin were available. To obtain such data, ^177^Lu-DOTA-knottin was synthesized, yielding a product with 98% radiochemical purity and a molar activity of approximately 44 GBq/µmol. The tracer was injected intravenously in mice bearing Capan2 xenografts on both shoulders. SPECT/CT images were taken at different times post injection. Figure 3A shows SPECT/CT images taken at 22 hours p.i. The maximum intensity projection (MIP) reveals an accumulation of ^177^Lu-DOTA-knottin in both the tumors and the kidneys. Uptake by other organs was low or moderate. Short-term and long-term tracer kinetics for kidney and tumor were quantified from SPECT images, up to 3 and 187 hours post-injection, respectively. Indeed, kidneys showed a higher initial accumulation of ^177^Lu-DOTA-knottin (Fig. 3B), yet clearance was also faster than from the tumor (Fig. 3C).

**Figure 3:**
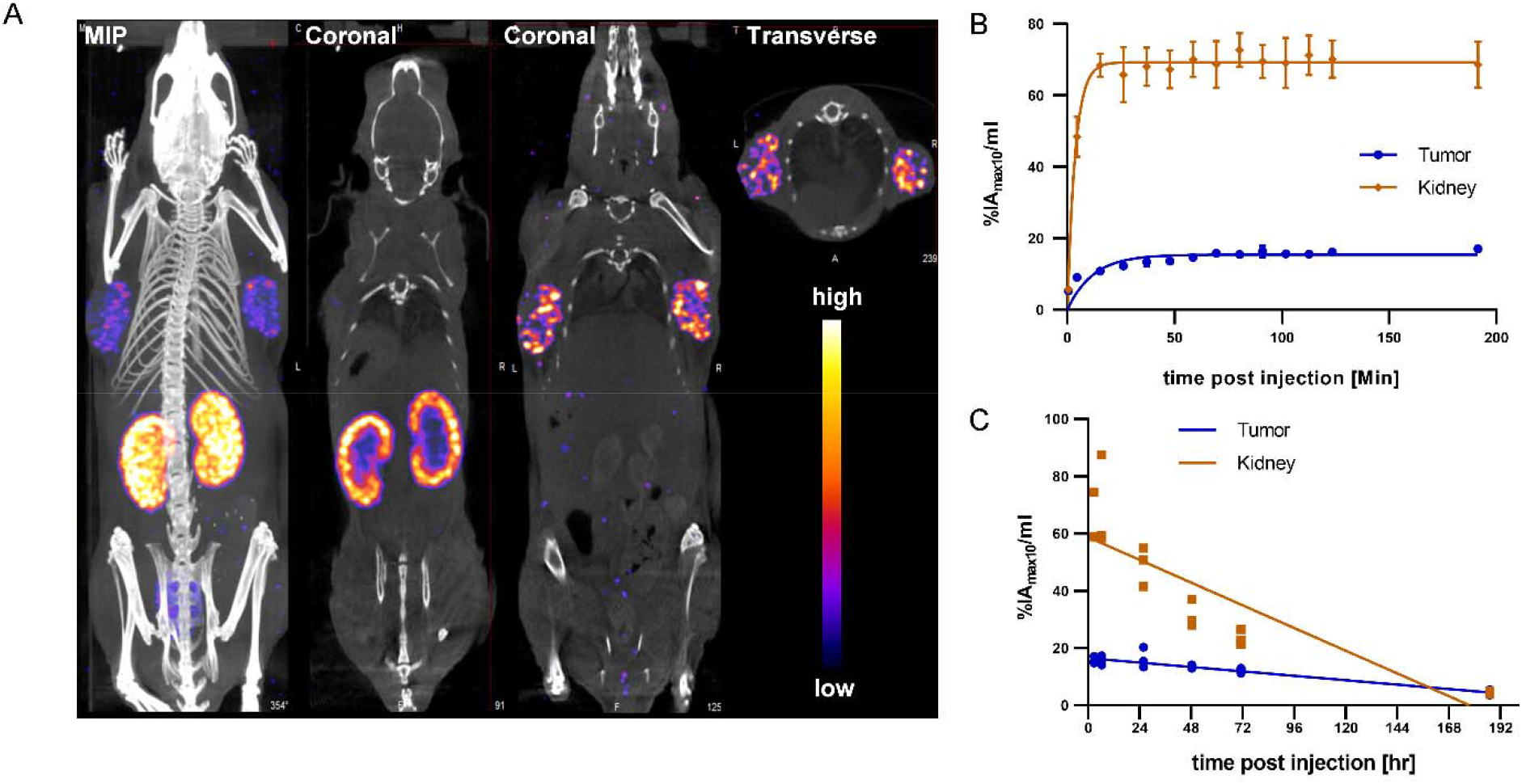
SPECT/CT imaging with ^177^Lu-DOTA-knottin. (A) SPECT images show maximum intensity projection (MIP), coronal and transverse projections fused with CT images of ^177^Lu-DOTA-knottin in a Capan2 xenograft model at 22 hours post injection of 62 MBq tracer. Nude mice were carrying xenografts on left and right shoulder. (B) Early SPECT kinetics data show the uptake of ^177^Lu-DOTA-knottin in tumor and kidney. (C) SPECT-based time-activity curve (2-187 hours p.i.) shows faster clearance of ^177^Lu-DOTA-knottin from kidney than from tumor.

### Biodistribution analysis of organ uptake

*Ex vivo* biodistribution analysis of mice bearing Capan2 xenografts demonstrated a tumor uptake of the ^177^Lu-DOTA-knottin of 3.1 ± 0.6, 2.5 ± 0.4, 3.5 ± 0.9 and 1.2 ± 0.2 %IA/g (mean ± S.E.M.) on day 1, 2, 3 and 8, respectively (Figure 4A). Nevertheless, tracer uptake by the kidney of 19.2 ± 2.5, 12.5 ± 0.6, 14.7 ± 4.5 and 2.3 ± 0.4 %IA/g was detected on days 1, 2, 3 and 8 p.i., respectively. On the other hand, low or moderate activity was discovered in organs like stomach, colon and lung at later time points (Table 2). The time-activity curve from *ex vivo* biodistribution data (Figure 4B) confirms the analysis of SPECT kinetics (Figure 3C) with activity from kidneys being washed out more rapidly than from tumors. Based on *ex vivo* biodistribution inputs, tumor-to-organ ratios were calculated (Table 2, Figure 5). Except for kidney, these ratios were favorable in pancreas, liver, blood, lung and muscle. Ex-vivo analysis of H&E-stained xenografts did not reveal significant differences in tissue structure between samples obtained on day 1 or day 8 after injection of the tracer (Supplementary Fig. 1).

**Figure 4:**
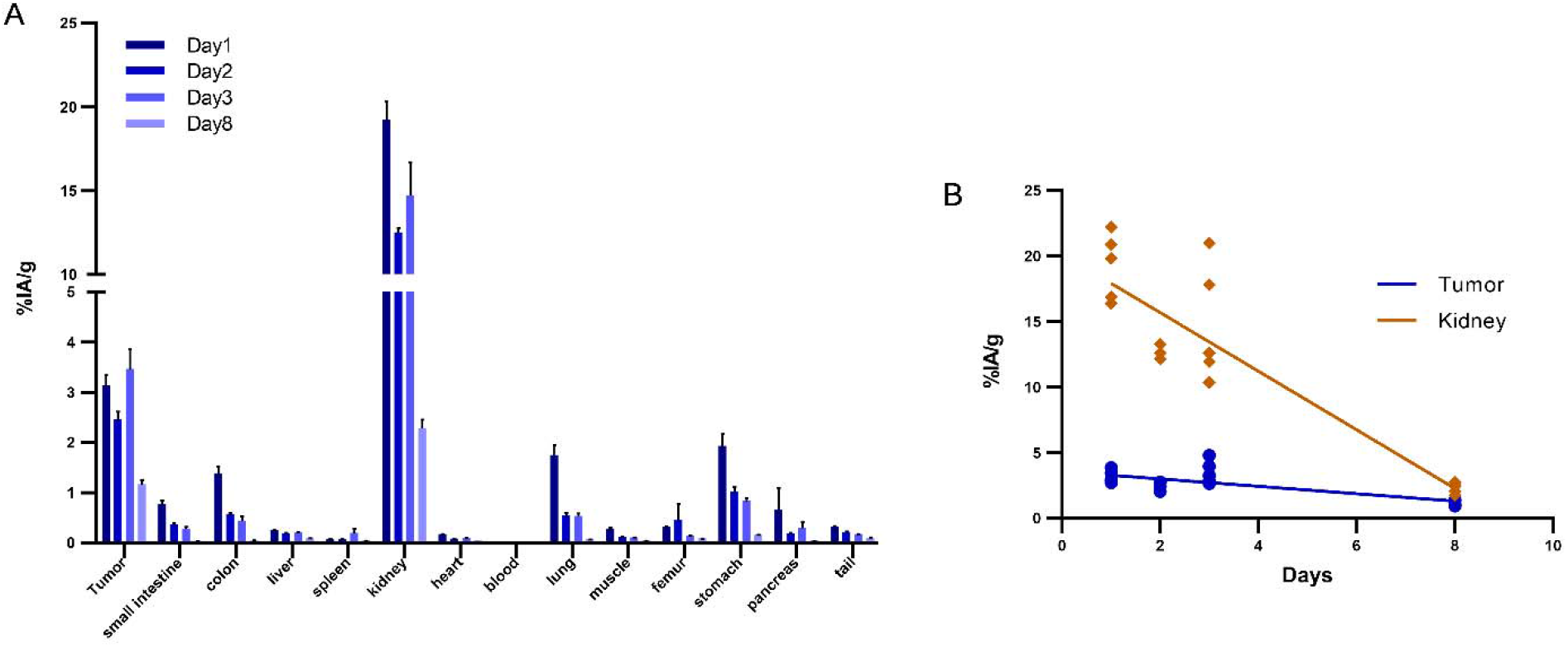
Biodistribution of ^177^Lu-DOTA-integrin-αvβ6 knottin in xenograft-bearing mouse. (A) Quantitative analysis of organ activity at day 1, 2, 3 and 8 p.i. (B) Time-activity curve of ^177^Lu-DOTA integrin αvβ6 knottin biodistribution data for tumor and kidney from days 1,2, 3 and 8 p.i.. Data represent mean ± S.E.M. (4 ≤ n ≤ 5).

**Table 2:** Biodistribution (%IA/g, mean ± SD) and tumor-to-organ ratios of ^177^Lu-DOTA-knottin in nude mice bearing Capan2 xenografts. Data are presented as mean ± SD %IA/g of tissue (4 ≤ n ≤ 5).

|  | 24 h |  | 48 h |  | 72 h |  | 192 h |  |
| --- | --- | --- | --- | --- | --- | --- | --- | --- |
| tumor | 3.13 | $\pm 0.63$ | 2.45 | $\pm 0.39$ | 3.46 | $\pm 0.85$ | 1.16 | $\pm 0.21$ |
| small intestine | 0.76 | $\pm 0.17$ | 0.35 | $\pm 0.08$ | 0.27 | $\pm 0.10$ | 0.02 | $\pm 0.00$ |
| colon | 1.37 | $\pm 0.36$ | 0.56 | $\pm 0.06$ | 0.43 | $\pm 0.21$ | 0.04 | $\pm 0.01$ |
| liver | 0.24 | $\pm 0.03$ | 0.18 | $\pm 0.04$ | 0.20 | $\pm 0.04$ | 0.08 | $\pm 0.02$ |
| spleen | 0.06 | $\pm 0.01$ | 0.07 | $\pm 0.01$ | 0.18 | $\pm 0.21$ | 0.04 | $\pm 0.01$ |
| kidney | 19.21 | $\pm 2.53$ | 12.50 | $\pm 0.55$ | 14.71 | $\pm 4.48$ | 2.28 | $\pm 0.40$ |
| heart | 0.16 | $\pm 0.03$ | 0.07 | $\pm 0.01$ | 0.08 | $\pm 0.04$ | 0.04 | $\pm 0.01$ |
| blood | 0.01 | $\pm 0.00$ | 0.00 | $\pm 0.00$ | 0.01 | $\pm 0.00$ | 0.00 | $\pm 0.00$ |
| lung | 1.74 | $\pm 0.46$ | 0.53 | $\pm 0.13$ | 0.52 | $\pm 0.13$ | 0.06 | $\pm 0.01$ |
| muscle | 0.26 | $\pm 0.09$ | 0.11 | $\pm 0.02$ | 0.10 | $\pm 0.02$ | 0.04 | $\pm 0.01$ |
| femur | 0.30 | $\pm 0.04$ | 0.45 | $\pm 0.65$ | 0.13 | $\pm 0.03$ | 0.07 | $\pm 0.02$ |
| stomach | 1.92 | $\pm 0.55$ | 1.01 | $\pm 0.19$ | 0.83 | $\pm 0.13$ | 0.14 | $\pm 0.03$ |
| pancreas | 0.65 | $\pm 0.98$ | 0.17 | $\pm 0.05$ | 0.29 | $\pm 0.26$ | 0.03 | $\pm 0.01$ |
| tail | 0.31 | $\pm 0.05$ | 0.20 | $\pm 0.05$ | 0.16 | $\pm 0.03$ | 0.08 | $\pm 0.05$ |
| tumor-to-blood | 497.70 | $\pm 79.30$ | 851.38 | $\pm 266.49$ | 697.02 | $\pm 248.45$ | 2123.36 | $\pm 553.90$ |
| tumor-to-kidney | 0.17 | $\pm 0.05$ | 0.19 | $\pm 0.04$ | 0.24 | $\pm 0.02$ | 0.51 | $\pm 0.07$ |
| tumor-to-muscle | 13.13 | $\pm 5.06$ | 23.15 | $\pm 4.30$ | 35.80 | $\pm 3.35$ | 32.86 | $\pm 13.74$ |
| tumor-to-liver | 12.99 | $\pm 1.72$ | 13.58 | $\pm 2.66$ | 18.35 | $\pm 6.29$ | 14.64 | $\pm 1.79$ |
| tumor-to-pancreas | 13.07 | $\pm 7.99$ | 15.56 | $\pm 7.69$ | 26.96 | $\pm 22.55$ | 38.69 | $\pm 10.34$ |

**Figure 5:**
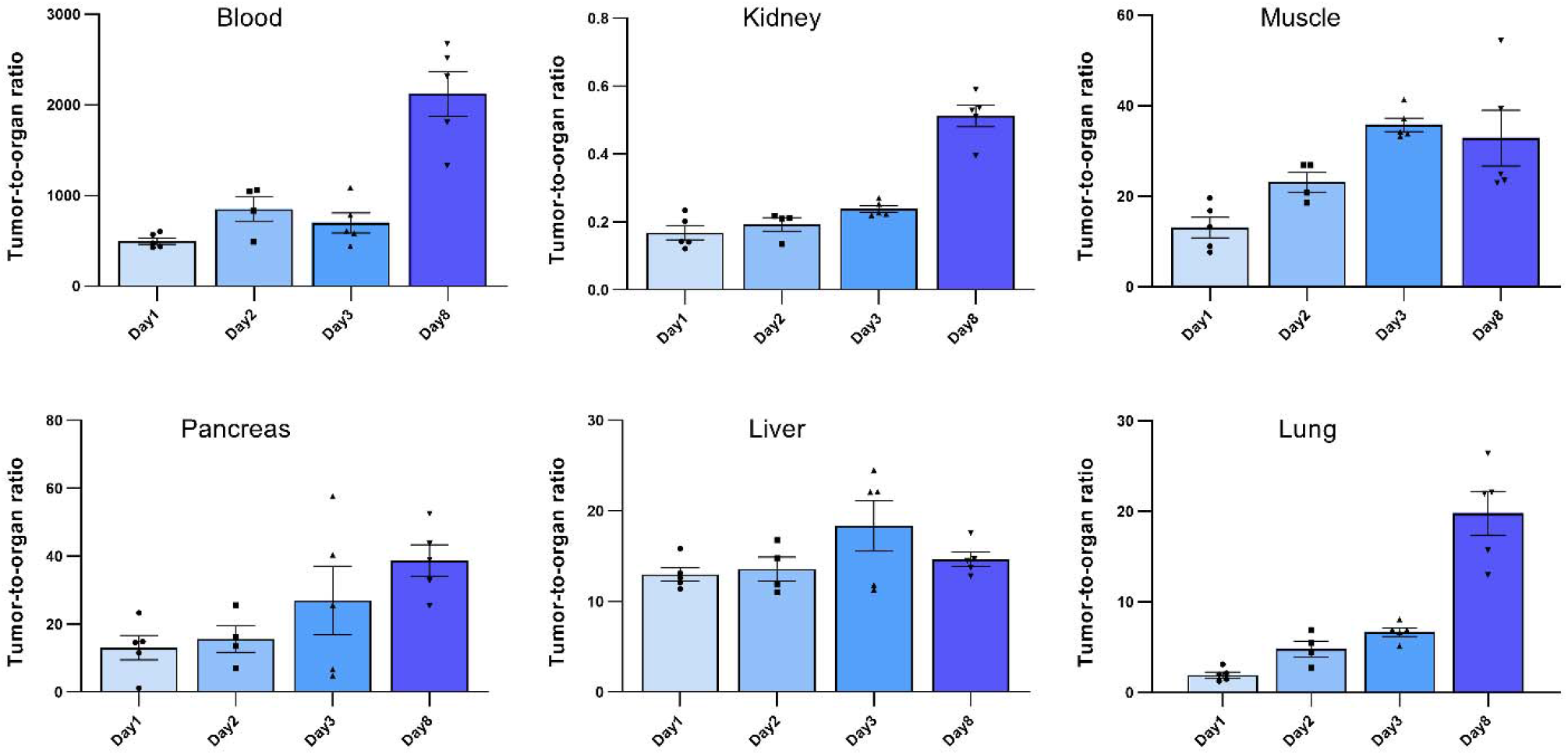
Tumor-to-organ ratios of the ^177^Lu-DOTA-integrin-αvβ6 knottin. Ratios to blood, kidney, muscle, pancreas, liver and lung were calculated using the *ex vivo* biodistribution values. Data represent mean ± S.E.M. (4 ≤ n ≤ 5).

### Dosimetric calculation

Dosimetric calculations for the human male adult for ^177^Lu ware generated (Table 3). The expected absorbed doses per injected activity in humans were calculated using the mouse biodistribution data. The two interspecies scaling (mass and time) model (*25,26*) was implemented for extrapolating animal to human data. The calculated expected effective dose was 0.04 mSv/MBq. Additionally, the calculated absorbed doses for the common organs are indicated. As expected from the mouse biodistribution data, kidney showed the highest absorbed dose of 0.02 mSv/MBq followed by lung and stomach of 0.01 and 0.005 mSv/MBq respectively (Table 3).

**Table 3:** Expected absorbed doses of ^177^Lu-DOTA integrin αvβ6 knottin in humans.

| **Target Organ** | **mSv/MBq** | **rem/mCi** |
| --- | --- | --- |
| Adrenals | 6.78E-06 | 2.51E-05 |
| Brain | 3.42E-08 | 1.26E-07 |
| Breasts | 2.36E-05 | 8.71E-05 |
| Gallbladder Wall | 0.00E+00 | 0.00E+00 |
| LLI Wall | 7.21E-05 | 2.67E-04 |
| Small Intestine | 1.82E-04 | 6.74E-04 |
| Stomach Wall | 5.15E-03 | 1.90E-02 |
| ULI Wall | 4.30E-06 | 1.59E-05 |
| Heart Wall | 0.00E+00 | 0.00E+00 |
| Kidneys | 2.06E-02 | 7.62E-02 |
| Liver | 9.34E-04 | 3.46E-03 |
| Lungs | 1.58E-02 | 5.85E-02 |
| Muscle | 1.40E-06 | 5.17E-06 |
| Ovaries | 1.76E-04 | 6.51E-04 |
| Pancreas | 5.59E-06 | 2.07E-05 |
| Red Marrow | 1.00E-04 | 3.71E-04 |
| Osteogenic Cells | 8.15E-06 | 3.01E-05 |
| Skin | 2.22E-06 | 8.20E-06 |
| Spleen | 2.35E-05 | 8.69E-05 |
| Testes | 0.00E+00 | 0.00E+00 |
| Thymus | 1.37E-06 | 5.08E-06 |
| Thyroid | 7.36E-06 | 2.72E-05 |
| Urinary BI Wall | 1.09E-05 | 4.02E-05 |
| Uterus | 1.99E-06 | 7.36E-06 |
| **Effective dose** | **4.31E-02** | **1.60E-01** |

## Discussion

Due to its high and selective tumor overexpression, integrin αvβ6 is emerging as a target in cancer for nuclear imaging. Several tumor-targeting strategies based on integrin αvβ6 have been developed for either diagnostic or therapeutic purposes (*15,27*–*31*). To that end, ligands had been conjugated either with a radionuclide (^18^F, ^64^Cu, ^111^In or ^177^Lu) or an anti-cancer drug (e.g. tesirine) (*31*). The majority of approaches involved the use of linear peptides derived from foot-and-mouth disease virus, which show a high affinity to the receptor. However, due to reduced stability and specificity for the receptor, *in vivo* results were ambiguous. In a recent study, a cyclic radiotracer specific for integrin αvβ6 (^68^Ga-cycratide) was used in a pancreatic mouse model for PET imaging (*32*). The integrin αvβ6 knottin applied here has recently been used in a first-in-human clinical study and has demonstrated a high potential for PDAC targeting (*24*). In this study, a DOTA-conjugated engineered cystine knot peptide (knottin) specific for integrin αvβ6 was chosen. The knottin peptide’s stability and affinity to the receptor were not compromised *in vivo* and it had previously exhibited favorable tumor uptake (*21,23*). Likewise, fluorescence-labeled knottins specific to avb6 had shown promising preclinical results, which could be further translated for the early detection of PDAC in patients (*33,34*). Such an optical imaging agent could also play a role in fluorescence-guided surgery for PDAC patients. Indeed, the specificity of the agent would assist in discerning PDAC from pancreatitis and normal pancreatic tissue.

In the current study, integrin αvβ6 mRNA expression in BxPC3 and Capan2 was found to be highest among all tested PDAC and other cell lines. Additionally, sustained expression of integrin αvβ6 in mouse xenografts of these cell lines was confirmed. More importantly, mRNA expression of integrin αvβ6 in mouse liver and kidney was found to be very low compared to BxPC3 and Capan2 cell lines. This, however, does not rule out expression in specific anatomical substructures of these two organs of excretion. Immunofluorescence experiments established the expression of integrin αvβ6 on the surface of Capan2 cells. This is of relevance as RT-qPCR data will only be informative about mRNA levels, not protein. In addition, many cell membrane receptors occur in an equilibrium of distribution between plasma membrane and intracellular compartments, e.g. the trans-Golgi network or endosomes. Proof of a high degree of surface expression may therefore be a meaningful predictor of in vitro and in vivo tracer binding.

Along with mRNA and protein expression data, the presence of a substantial number of functional receptors on the cell surface is an essential benchmark for nuclear imaging. In agreement with previous findings, the affinity of the knottin for BxPC3 and Capan2 cells was found to be in the low nanomolar range (1.69 and 9.46 nM) as revealed by radioligand binding assay. It is interesting to note that the affinity of the knottin did not change upon either the addition of a DOTA chelator moiety or incorporation of natural non-radioactive lutetium (^nat^Lu) into the chelator.

Compared to previous biodistribution studies (^18^F, ^99m^Tc) using the same knottin (*36,37*), a higher uptake of the ^177^Lu tracer in tumors was observed here. Some organs like lung, stomach, colon and kidney showed tracer uptake at early time points yet this was washed away at later time points. As integrin αvβ6 expression in mouse kidney had been found several orders of magnitude lower than in xenograft tumors (Fig. 1A), this uptake in the renal medulla may be attributed to unspecific uptake by transporters, e.g. OATPs. A high amount of tracer was accumulated in the kidney at early time points (day 1: 19.2 ± 2.5 %IA/g); there was a significant reduction at later time points (day 8: 2.3 ± 0.40 %IA/g). However, in the tumor the activity was cleared at a much lower rate than in the kidney. Higher tracer activity in the kidney resulted in a lower tumor-to-kidney ratio at day-1, however, Capan2 tumors were clearly recognized. Similarly, a higher tumor-to-organ ratio at all-time points gives an advantage for contrast and favorable tumor imaging. These findings again underline that the tracer binds in the tumor specifically and with greater affinity. However, tumor activity is still limiting the full impact of this targeting approach. A further improved initial tracer uptake in the tumors should be the goal for the continued path towards translation into the clinic. With affinities already in a very favorable range, overall design of the conjugate could be modulated, e.g. by the introduction of a different spacer. In addition, increasing the molar activity of the tracer formulation may result in increased tumor uptake.

For dosimetry, biodistribution data were extrapolated to human adult male using OLINDA/Exm software for the therapeutic radionuclide ^177^Lu. This dosimetric projection provided a preliminary estimate for assessing the therapy-associated risk of radiation damage. The absorbed doses in the kidney propose it as the dose-limiting organ. As this method involves two scaling methods to extrapolate human data, the resulting prediction is of preliminary nature and needs further confirmation, e.g. by collecting corresponding data from human subjects.

In the framework of this study, only a single dose of tracer was injected into the mice, and the animals were monitored for no longer than eight days. Consequently, no significant impact was observed in H&E-stained tissue sections of day 1 and day 8 post injection. A therapy study, potentially involving multiple/repeated administrations of tracer is required to establish therapeutic efficacy. Still, the uptake of tracer by xenograft tumors at different time points and faster renal clearance confirms the suitability of the knottin peptide as a tracer and of integrin αvβ6 as a valid target for diagnosis and potential targeted radionuclide therapy in PDAC.

## Conclusion

In this study, binding of the peptide tracer ^177^Lu-DOTA-integrin αvβ6 knottin to its target both *in vitro* and *in vivo* was investigated. ^177^Lu-DOTA-integrin αvβ6 exhibited high affinity and specific binding to target-positive cells and tumors. The study demonstrated the translational potential of this tracer for imaging and therapy of integrin αvβ6-overexpressing tumors like PDAC.

## Abbreviations

% IA/g: percent injected activity per gram tissue
^177^Lu: lutetium-177
BERIC: Berlin Experimental Radionuclide Imaging Center
BSA: bovine serum albumin
CPM: counts per minute
DMSO: dimethyl sulfoxide
DOTA: 1,4,7,10-tetraazacyclododecane-1,4,7,10-tetraacetic acid
EC_50_: half maximal effective concentration
ECM: extracellular matrix
HPLC: high-performance liquid chromatography
IC_50_: half maximal inhibitory concentration
n.d.: not determined
NET: neuroendocrine tumor
NMRI: Naval Medical Research Institute
OATP: organic anion transporter protein
p.i.: post injection
PBS: phosphate-buffered saline
PDAC: pancreatic ductal adenocarcinoma
PFA: paraformaldehyde
PET: positron emission tomography
PRRT: peptide receptor radionuclide therapy
RPMI1640: Roswell Park Memorial Institute medium 1640
SD: standard deviation
SEM: standard error of mean
SSA: somatostatin analog
SSTR: somatostatin receptor
TE: echo time
TFA: trifluoroacetic acid

## Author Contributions

Concept and experimental design: BW, SSG, RK, SR, CG. Development of methodology: RK, NB Acquisition of data: SS, TH, SE, SP, NB. Analysis and interpretation of data: SS, CG. Writing, review, and/or revision of the manuscript: SS, CG and all other authors. Study supervision: BW, SSG, CG.

## Acknowledgements

We thank Ines Eichhorn for expert technical assistance.

