## Supplemental data for "SPECT/CT imaging, biodistribution and radiation dosimetry of a ^177^Lu-DOTA-integrin αvβ6 cystine knot peptide in a pancreatic cancer xenograft model"

**Supplementary Figures**

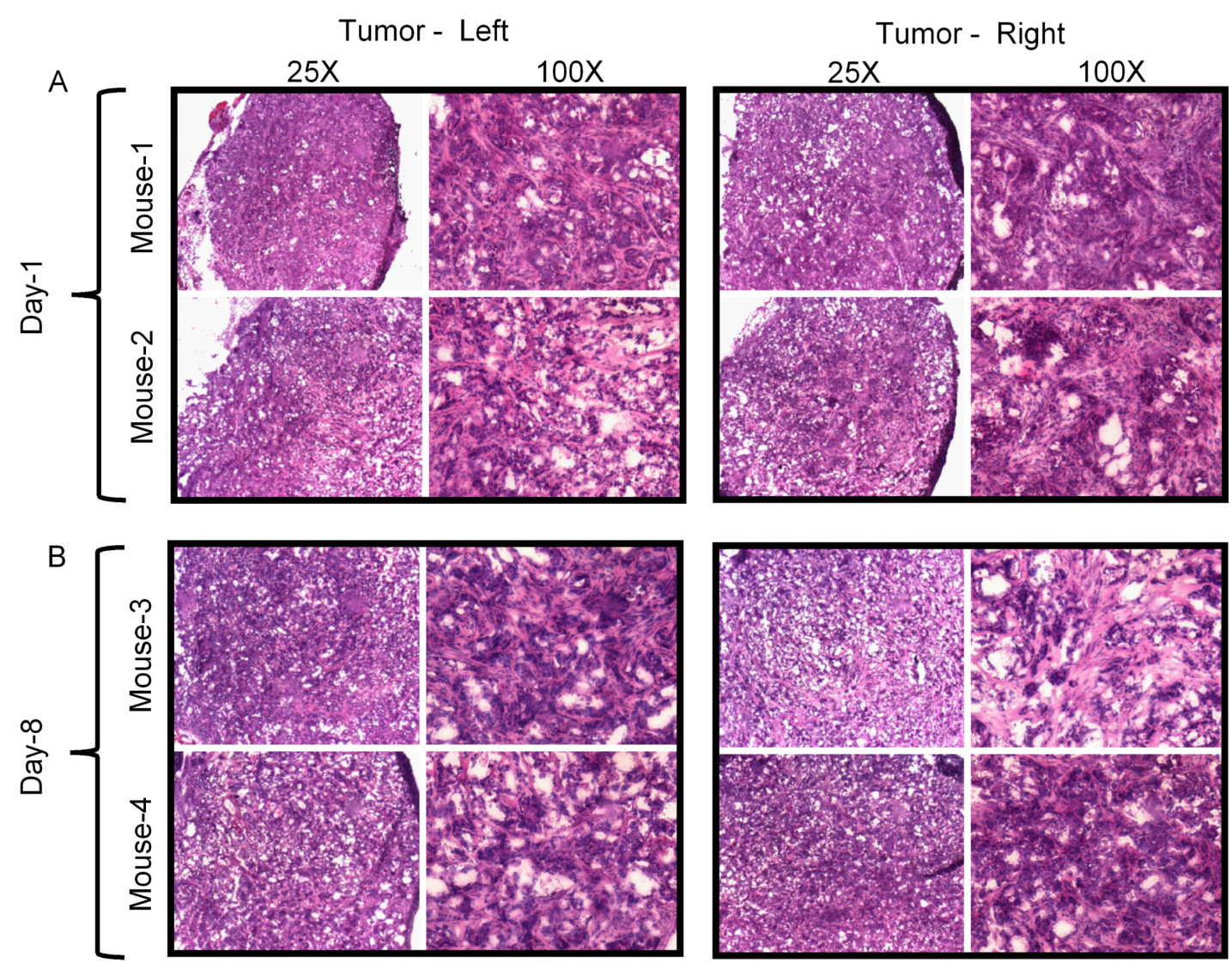
**Supplementary Figure S1**

**Supplementary Figure S1: H&E staining of tumor tissue**. Comparison of H&E staining of the tumor (left and right) tissue from different mice at day-1 and day-8 post-injection
